# Different contrast encoding in ON and OFF visual pathways

**DOI:** 10.1101/2020.11.25.398230

**Authors:** Saad Idrees, Thomas A. Münch

## Abstract

Subjective visual experience builds on sensory encoding of light reflected by different objects in our environment. Most retinal ganglion cells encode changes in light intensity, quantified as contrast, rather than the absolute intensity. Mathematically, contrast is often defined as a relative change in light intensity. Activity in the visual system and perceptual responses are usually explained with such definitions of contrast. Here, for the first time, we explicitly explored how contrast is actually represented in the visual system. Using mouse retina electrophysiology, we show that response strength of OFF retinal ganglion cells does not represent relative, but absolute changes in light intensity. ON RGC response strength is governed by a combination of absolute and relative change in light intensity. This is true for a wide range of ambient light levels, at least from scotopic to high mesopic regimes. Consequently, light decrements and increments are represented asymmetrically in the retina, which may explain the asymmetries in responses to negative and positive contrast observed throughout the visual system. These findings may help to more thoroughly design and interpret vision science studies where responses are driven by contrast of the visual stimuli.

## Introduction

The activity of retinal ganglion cells (RGCs) does not encode the absolute light intensity, with the notable exception of intrinsically photosensitive retinal ganglion cells (ipRGCs), also known as melanopsin-containing ganglion cells (Do et al., 2009). Instead, ganglion cells encode the *change* in light intensity. The magnitude of this change is called contrast. In this study, we asked two questions: according to which rules do retinal ganglion cells encode contrast? And is this interpretation of contrast different at different light intensity regimes, such as for night-time vision and day-time vision?

Mathematically, contrast can be quantified in different ways, which we will exemplify with a real-world scenario. Imagine the sun breaking through the clouds; the world becomes brighter by a certain amount. How big, quantitatively, is this intensity increase? Let us consider different objects in the scene that we are looking at, for example a relatively bright flower (F) and a relatively dark leaf (L, compare inset in Fig. 3). “Bright” and “dark” here mean that these objects reflect different amounts of the incident light. Thus, while the sun has still been behind the clouds, those objects had different starting intensities (*F_Cloud_* and *L_Cloud_*), and after the “event” they have different end intensities (*F_Sun_* and *L_Sun_*). We can assume that the reflectance of the flower and leaf are fixed physical properties and do not change (Land and McCann, 1971; Shapley and Enroth-Cugell, 1984), so that the intensity changes by a constant and identical factor *k* > 1 when the sun breaks through the clouds, namely *Intensity_after_* = *k* · *Intensity_before_*. We then get *F_sun_* = *k* · *F_Cloud_* and *L_sun_* = *k* · *L_Cloud_*, or 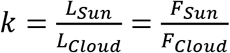. This ratio is one way of expressing how much the intensities of objects have changed:

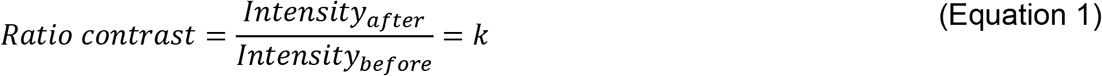

Weber and Michelson contrasts are other ways to express the change in intensity. If, again, we assume that *Intensity_after_* = *k Intensity_before_*, we get

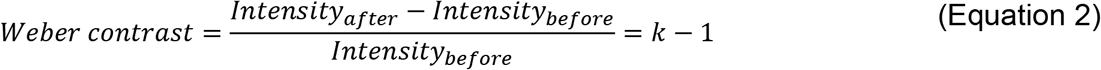

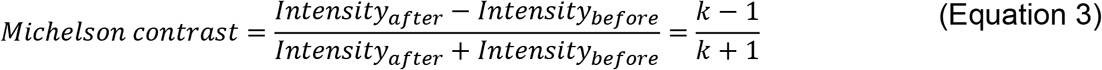

Numerically, Ratio contrast, Weber contrast, and Michelson contrast give different values, but they have in common that the contrast value only depends only on *k* and is independent of the absolute intensity of objects in the world. In other words: the value is the same for the flower as for the leaf. As such, these three different measures of contrast express relative changes in light intensity that do not depend on the initial intensity.

One could also quantify the change in light intensity differently, for example by how much (in absolute terms) an object’s intensity has increased. Under the same assumption as before, namely *Intensity_after_* = *k* · *Intensity_before_*, we get

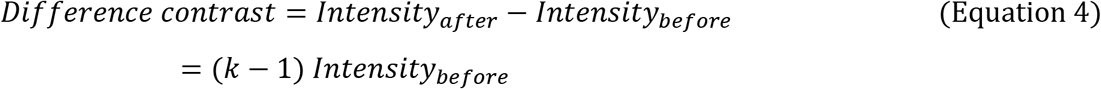

In this metric, the brightness change of each object is not independent of its initial intensity, but it changes by the factor of *k* – 1 of its initial brightness. Here, the bright flower and the dim leaf produce a different contrast when the sun breaks through the clouds. Thus, “Difference contrast” provides a fundamentally different interpretation of the event as Ratio contrast, Weber contrast, or Michelson contrast.

Figure 1a and b depict the different scenarios described above. In both plots, the x-axis represents the initial intensities of objects, the y-axis the intensities after light change. Any point in this coordinate system therefore corresponds to an *Intensity_before_* → *Intensity_after_* event that has a certain contrast. If two points fall on the same line in Fig. 1a, then they have the same contrast according to the interpretation provided by Ratio contrast (and also by Weber or Michelson contrast): gray lines represent increases in light intensity (*k* > 1, which is equivalent to (*k* – 1) > 0, or 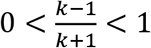), and different gray lines represent events where light intensity increases by different amounts. Black lines, correspondingly, show various iso-contrast conditions for decreases in light intensity (0 < *k* < 1). In Fig. 1b, the lines indicate iso-contrast events according to the interpretation provided by Difference contrast. Note that the iso-contrast lines in Fig. 1a are increasingly steep, while they are parallel to each other in Fig. 1b.

**Figure 1.**
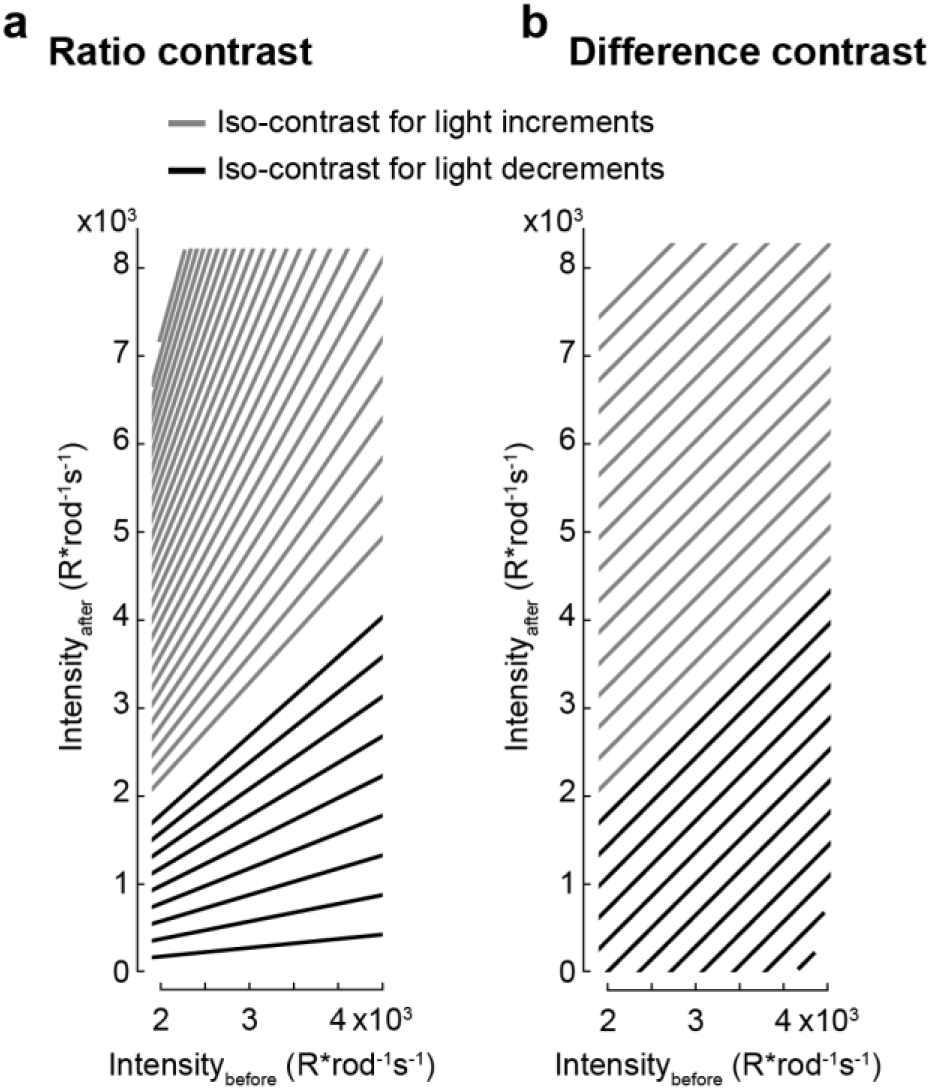
Iso-contrast lines for different before/after intensity combinations. The x- and y-axes represents the intensity (in R*rod^-1^s^-1^) of objects before and after experiencing luminance change, respectively. Contrast can be calculated for each *Intensity_before_* → *Intensity_after_* combination, for example in the forms given by Equations 1–4. *Intensity_before_* → *Intensity_after_* combinations that fall on any one line have the same contrast according to Equations 1–3 (**a**) or Equation 4 (**b**). Gray lines: increase in intensity (Intensity_after_ > Intensity_before_). Black lines: decrease in intensity (Intensity_after_ < Intensity_before_).

How is contrast represented in the responses of retinal ganglion cells (RGCs)? Our hypothesis was that RGCs would encode relative changes in light intensity, i.e. their response strength would be proportional to the Ratio, Weber, or Michelson contrast experienced by them. If true, an ON RGC would then treat the intensity change of the flower (*F_Cloud_* → *F_Sun_*) and the leaf (*L_Cloud_* → *L_Sun_*) as equivalent and would respond with comparable strength to those two events. Correspondingly, an OFF RGC would respond to the opposite event, when the sun hides behind clouds, in the same way, irrespective of the object that it is exposed to (flower or leaf). Independent of the validity of our hypothesis, it is clear that an RGC may respond with the same strength to different stimulus events, i.e. different combinations of before/after intensities. Our approach has been to record the responses of RGCs to many such before/after combinations, and draw “iso-response lines” for RGCs similar to the “iso-contrast lines” in Fig. 1. If our hypothesis is true, namely that RGCs faithfully encode light changes as the ratio of before/after light intensities, then we would expect that these iso-response lines follow the same trend as the iso-contrast lines of Fig. 1a. Otherwise, in contradiction to our hypothesis, we may observe other scenarios, such as iso-response lines that are parallel to each other (Fig. 1b, representing the scenario that the “Difference Contrast” is the relevant metric for RGCs, i.e. they would encode absolute change, rather than relative change of light intensity), or that the iso-response lines become increasingly less-steep (representing a scenario with response suppression, for example when responses start to become saturated). However, in our experiments, we tried to avoid this last scenario by restricting stimuli to moderate contrasts. We found that the behavior of ON RGCs was consistent with our hypothesis, they appear to encode relative changes in light intensity. OFF RGC behavior, on the other hand, was inconsistent with the hypothesis, and they seem to encode absolute changes of light intensities.

## Results

To test the hypothesis that responses of RGCs solely depend on the relative change in light intensity irrespective of the starting intensity, we recorded the spiking activity of RGCs in isolated ex vivo mouse retinae (n=3) using high-density multi-electrode arrays (Müller et al., 2015) (MEAs), while we exposed them to several different step stimuli. Each step stimulus consisted of a uniform background, of one of 16 possible intensities, that was presented for 4 seconds. We refer to this intensity as *Intensity_before_*. Then, the intensity was increased or decreased instantaneously (“ON step” or “OFF step”) and stayed at the new value (*Intensity_after_*) for 1 s. In total, there were 368 ON steps (combinations of before and after intensities), and 256 OFF steps, repeated several times. Most RGCs responded robustly to these steps. We quantified the response strength as the peak of the RGC’s spike rate within 400 ms after the step. Different *Intensity_before_* → *Intensity_after_* steps resulted in different response strengths. To quantify the response at the population level, we first normalized the responses of each recorded RGC relative to its median response strength to all *Intensity_before_* → *Intensity_after_* combinations (analyzing ON steps for the n=177 ON RGCs; analyzing OFF steps for the n=66 OFF RGCs. We normalized each cell by its median response strength, rather than by the maximal response, so that potential saturation of responses for stronger stimuli, or any outlier responses, would not influence the overall shape of the stimulus-response relationship.) We then generated a generic ON RGC by taking the median response strengths to each before/after combination across all ON cells. Finally, we normalized the resulting responses (Fig. 2a, black dots). We fitted a second-order polynomial to these responses to estimate the response strength of the generic RGC to a continuous range of *Intensity_before_* → *Intensity_after_* combinations (Fig. 2a, surface fit). Similarly, we generated a generic OFF RGC (Fig. 2a).

**Figure 2.**
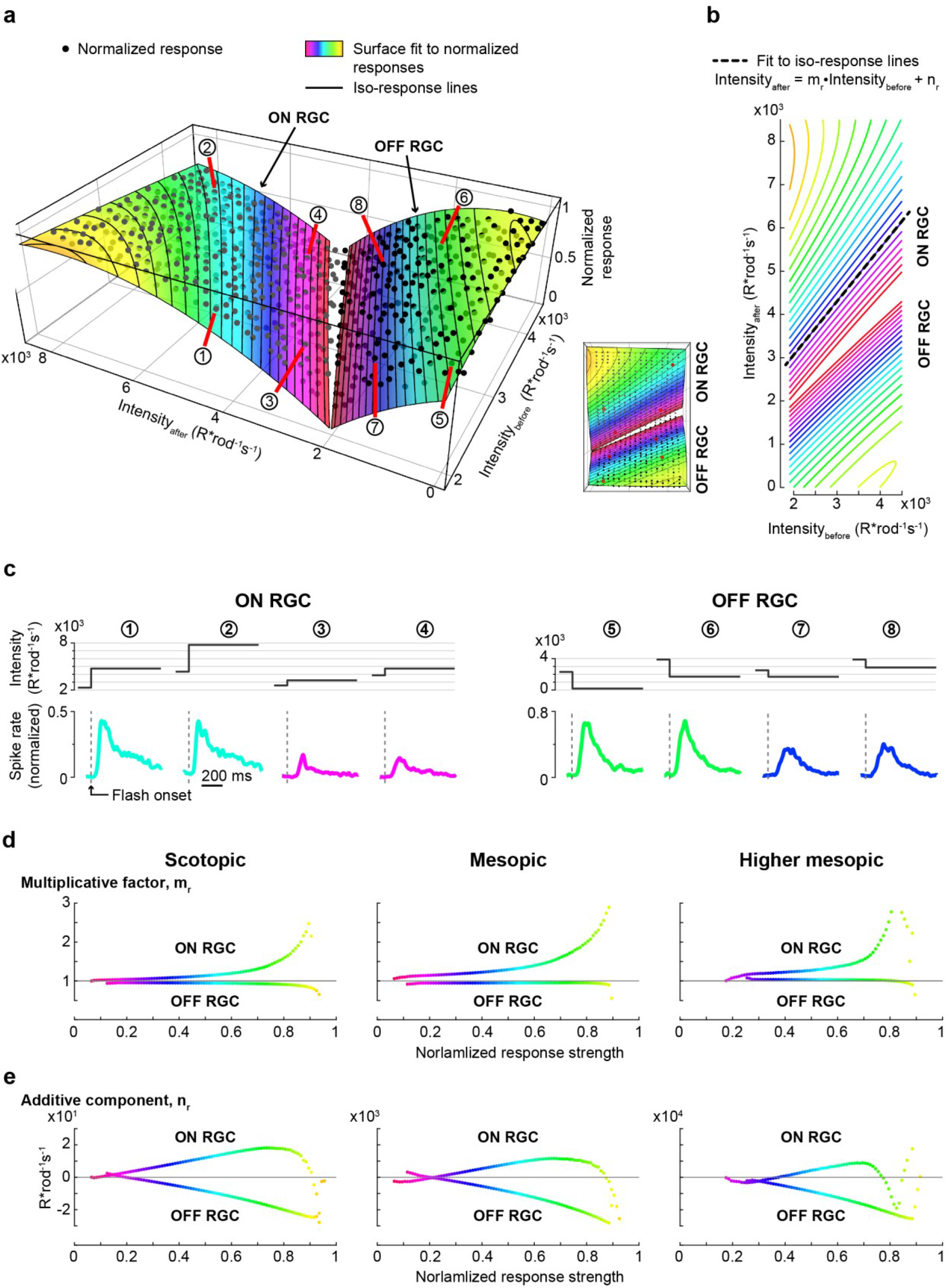
Generic RGC responses to Intensity_before_ → Intensity_after_ steps. **a. Left:** Normalized responses of generic ON and OFF RGCs as a function of *Intensity_before_* and *Intensity_after_* at mesopic ambient light levels. Intensities are reported in R*rod^-1^s^-1^. Dots correspond to the *Intensity_before_ → Intensity_after_* steps at which data was recorded (N = 368 steps with Intensity_after_ > Intensity_before_ that were used to analyze ON RGCs, and N = 256 steps with Intensity_after_ < Intensity_before_ for OFF RGC). The two surfaces were obtained by separately fitting a two-dimensional second-order polynomial to the ON RGC data and OFF RGC data. Different *Intensity_before_* → *Intensity_after_* steps that induced the same response strength can be found along iso-elevation lines of the surface, indicated by the same surface coloring, and highlighted by the black elevation lines. Circled numbers point to individual *Intensity_before_ → Intensity_after_* steps (1-4 for the generic ON RGC and 5-8 for the generic OFF RGC), for which the responses are plotted in **c. Right:** top view of the data shown on the left. **b.** Iso-response lines of the surface in **a** in a 2-D view, similar to the format of Fig. 1. Lines are color-coded to represent the same response strengths as in **a**. Black dashed line is the linear fit to one of these iso-response lines according to (Equation *5*). **c.** Normalized spike rate of the generic ON and OFF RGCs to the different *Intensity_before_ → Intensity_after_* steps indicated by circled numbers in **a**. (ON RGC: 1-4; OFF RGC: 5-8). The stimulus is shown above the response traces. Dashed gray lines mark the times of intensity change. Response traces are color-coded according the response strength in **a**; response strength is defined as the peak of the response trace. **d,e.** Values of the multiplicative factor *m_r_* **(d)** and the additive component *n_r_* **(e)** as a function of response strength *r*, for the generic ON and generic OFF RGC. These values were estimated by fitting (Equation *5*) to iso-response lines at intervals of 0.01. An example for such a fit is shown as dashed black line in **b**. Columns correspond to different ambient luminance levels, Scotopic: 0.09 R*rod^-1^s^-1^ to 82 R*rod^-1^s^-1^; medium mesopic: 9 R*rod^-1^s^-1^ to 8169 R*rod^-1^s^-1^; high mesopic: 91 R*rod^-1^s^-1^ to 81,690 R*rod^-1^s^-1^. Data in **a-c** shows results at mesopic light conditions.

Different *Intensity_before_* → *Intensity_after_* steps that induced the same response strength can be found along iso-elevation lines of the surface, indicated by the same surface coloring, and highlighted by the black elevation lines in Fig. 2a. For example, even though the stimulus steps marked 1 and 2 in Fig. 2a have different *Intensity_before_* and *Intensity_after_* values, they elicit very similar responses (Fig. 2c). Correspondingly, stimulus steps marked 3,4 of the generic ON RGC, and steps 5,6 and 7,8 of the generic OFF RGC elicit very similar responses (Fig. 2c). Fig. 2b shows these surface iso-elevation lines in a 2-dimensional view, similar to the format of Fig. 1. For low to medium intensity stimuli, these iso-response lines can be captured well with a linear regression model of the form

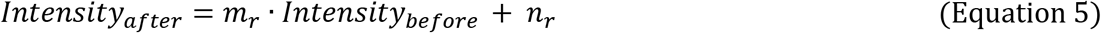

An example for such a linear fit is shown in Fig. 2b as a dashed black line. Figs. 2d and 2e show the parameters *m_r_* and *n_r_* of the linear regression as a function of response strength, *r*. For high-intensity stimuli the iso-elevation lines curve strongly (Fig. 2b); this corresponds to the surfaces in Fig. 2a flattening, meaning that the RGC responses start to saturate. For those high-intensity stimuli, a linear approximation of the iso-response lines according to (Equation *5* is not very meaningful. In Figs. 2d and 2e this becomes apparent for response strengths beyond 0.7. We will therefore limit our interpretation of the results to the range below 0.7, i.e. to the range of low to medium intensity stimuli.

We have performed the measurements and analysis described above at three light levels: scotopic, medium mesopic, and high mesopic. Fig. 2a-c shows the data for the medium mesopic light level, Fig. 2d-e show the parameters *m_r_* and *n_r_* for all three light levels.

At all light levels, we found that ON and OFF RGCs interpret “contrast” in different ways. For the generic OFF RGC, the multiplicative factor *m_r_* hardly varied with response strength *r* (Fig. 2d). This corresponds to the iso-response lines in Fig. 2b being parallel to each other (their slope, *m_r_*, does not change). With increasing response strength, these parallel lines move further down, represented by the ever-increasing negative values of the parameter *n_r_* (Fig. 2e). Note that this is the same situation as depicted in Fig. 1b, indicating that OFF RGC responses are almost exclusively driven by *absolute* changes in light intensities (*Intensity_after_* - *Intensity_before_*). In ON RGCs, on the other hand, the multiplicative factor *m_r_* rises continuously with response strength *r*, i.e. the iso-response lines in Fig. 2b have increasing slopes. This is similar to the situation depicted in Fig. 1a, indicating that ON RGCs are driven by relative changes in light intensity.

What are the implications of this OFF RGC behavior? We return to our example from the introduction. When the sun hides behind clouds, the world becomes darker, and objects (for example the flower and the leaf) all reflect less light by a factor of *k* (0 < *k* < 1). This real-world change of object intensities is represented in Fig. 3 by the two circles that are located on the black line with slope *k*. The gray dashed lines represent other scenarios, for example when the sun would be obscured by a lighter cloud leading to less darkening, or by a thicker cloud leading to more darkening. In the given situation, corresponding OFF RGCs exposed to the flower and the leaf will not respond equally to the event, because OFF RGCs follow the rules of difference-contrast. This is represented by the parallel red lines in Fig. 3: the flower and the leaf fall on different red lines; the brighter flower is located on an iso-response line corresponding to stronger responses than the iso-response line containing the leaf (compare Fig. 2a, the surface rises towards the bottom right). Correspondingly, the flower-OFF-RGC would respond more strongly to the event than the leaf-OFF-RGC because the absolute decrement is stronger, even though the illumination intensity for the two OFF RGCs decreases by the same factor. In general, when the world is dimming by a constant factor, OFF RGC responses to the dimming of brighter objects in the scene (flower) are stronger than responses to the corresponding dimming of darker objects (leaf).

**Figure 3.**
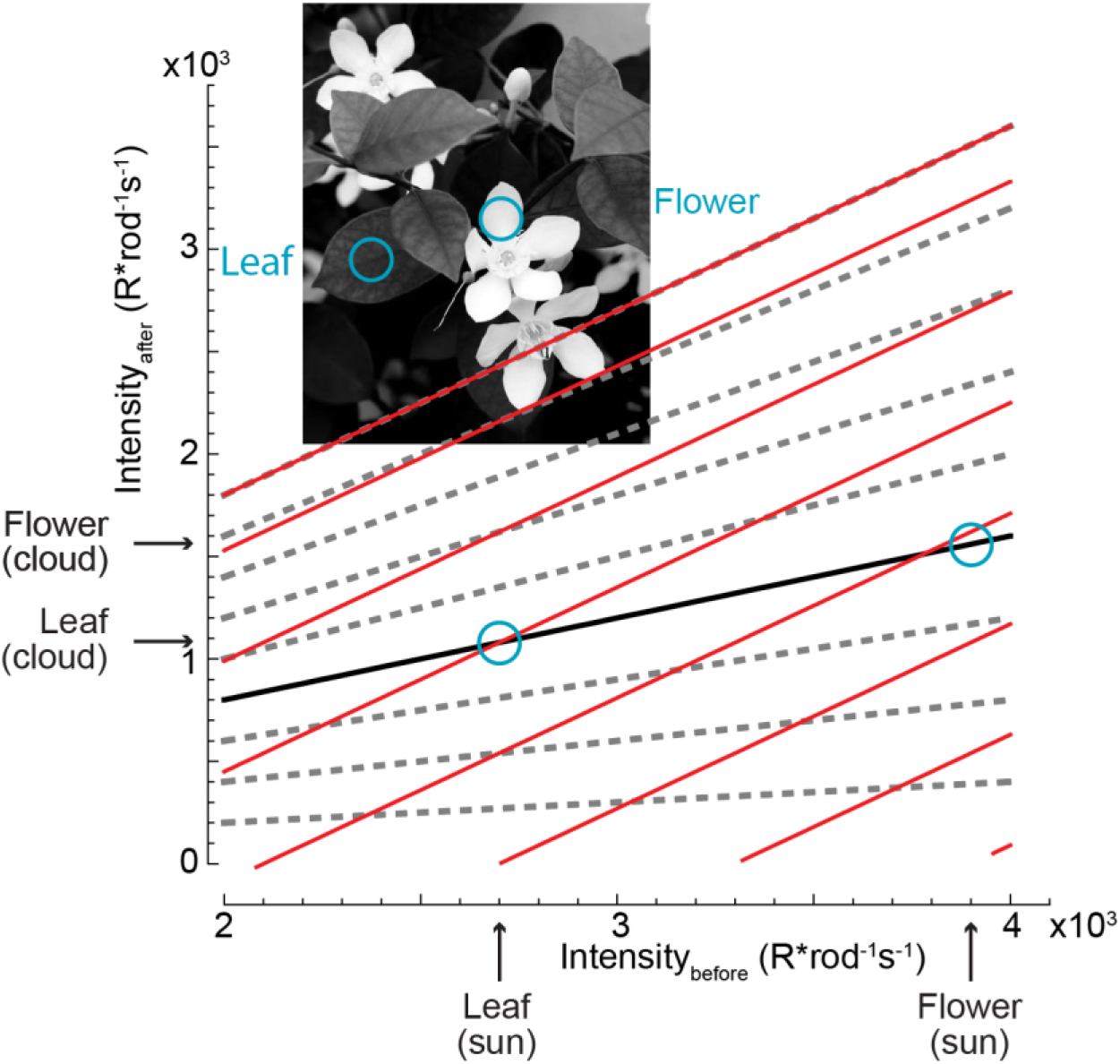
Implications of OFF RGC behavior according to Difference contrast. Image of a bright flower and a darker leaf illuminated by a single source (in this case light from the sun). The initial brightness of the hypothetical leaf and the flower in R*rod^-1^s^-1^, *Intensitiy_before_*, is marked by the arrows on x-axis. After a cloud covers the sun, their brightness decreases (*Intensitiy_after_*, y-axis). Thicker or thinner clouds would trigger a stronger or weaker off-stimulus, represented by the different dashed gray lines. Stronger off-stimulus corresponds to lower lines. Blue circles indicate the before/after intensities of the leaf and the flower for an example cloud that reduces the light falling on all objects by a factor of *k* (represented by black line; here k~2.5^-1^). Parallel red lines illustrate OFF RGC iso-responses that follow the difference-contrast. The flower and the leaf fall on different red lines; the brighter flower is located on an iso-response line corresponding to stronger responses.

When we inspect the behavior of ON RGCs more closely, we can observe that the additive parameter *n_r_* (Fig. 2e) is also not constant. Rather, *n_r_* appears to follow mirror-symmetric trends for ON and OFF RGCs. If ON RGCs would have behavior that purely adheres to Ratio contrast, the parameter *n_r_* should have a constant value of 0 (note that the hypothetical iso-contrast lines in Fig. 1a, as the dashed gray lines in Fig. 3, all intersect at the origin of the coordinate system: their intercept *n_r_* is always 0). This is clearly not the case for ON RGCs. So, like for OFF RGCs, ON RGC responses are enhanced for objects experiencing stronger absolute changes, i.e. darker objects in the scene. This mirrors OFF RGC behavior. However, given the fact that a component of ON RGC behavior is driven by Ratio contrast, this enhancement is not as pronounced as for OFF RGCs.

Taken together, at all three light intensities tested, a component of what drives RGC responses is the absolute change in light intensities, and this component of RGC behavior can be described by Difference contrast. In scenarios where the luminance of all objects changes by a constant factor, this emphasizes the responses of OFF RGCs to brighter objects, and of ON RGCs to darker objects. For OFF RGCs, this absolute brightness change appears to be the main, if not only, factor that determines their response strength. ON RGC responses are additionally driven by relative changes of brightness (according to Ratio, Weber, or Michelson contrast). Taken by itself, this would mean that ON RGCs would respond to all objects in a heterogeneous scene equally when the illumination of that scene increases by a constant factor. When combined with the behavior according to Difference contrast, this means that responses of different ON RGCs are more equalized than those of OFF RGCs.

## Discussion

Our results describe a novel asymmetry in ON and OFF RGCs with respect to what contrast means for them. For moderate changes in intensities, the peak spiking rate of OFF RGC responses represents absolute changes in intensities, in a manner consistent with Difference contrast. When we consider a scenario where the illumination of a scene changes by a constant factor, OFF RGC responses emphasize bright objects in a scene. ON RGCs, on the other hand, encode mostly relative changes in light intensity, in a manner consistent with the definition of Ratio, Weber, and Michelson contrast. While this would lead to equal responses to all object in a scene, responses of ON RGCs are also partially driven by absolute changes in light intensity, so that ON RGCs somewhat emphasize their responses to the brightening of dark objects.

ON and OFF RGCs are driven not only by global changes in illumination, but by any brightness change within their receptive fields. Arguably the most common scenario underlying such local brightness changes would be self-movement of the observer (eye and body movement) which leads to translational shifts of the projected world on the retina. These local changes in brightness are less coherent or predictable on the global scale as described for our original scenario. Still, the response rules we have discovered (how contrast is encoded differently by ON and OFF RGCs) are very fundamental and likely govern retinal activity also under these other scenarios.

The asymmetric representation of contrast in ON and OFF RGCs might be conserved along the visual hierarchy, contributing at least in part to the perceptual asymmetries in detecting light increments and decrements. For example, to generate equal psychophysical responses, light increments and decrements must be different in magnitude (Lu and Sperling, 2012), with decrements having a lower detectability threshold (Bowen et al., 1989; Lu and Sperling, 2012; Whittle, 1986).

What mechanisms in the retina may lead to the asymmetry in contrast representation across ON and OFF RGCs? Cone responses in various vertebrates have been shown to follow Weber’s law for moderate intensity changes (Burkhardt, 1994; Clark et al., 2013; Normann and Werblin, 1974; Shapley and Enroth-Cugell, 1984). Those observations were however mostly based on light increments. It is now well established that cone responses to light increments and decrements are asymmetric: the depolarization following a dark flash is larger in amplitude than the hyperpolarization following a bright flash of same magnitude from the background (defined as absolute difference) (Baden et al., 2013; Clark et al., 2013; Cooper, 2016). It is therefore likely that the mechanisms underlying different contrast representations in ON and OFF RGCs start already at the level of cones. This would mean that the underlying mechanism is independent of specialized circuitry. In addition, because ON and OFF RGCs show equally different contrast encoding at scotopic light levels (Fig. 2d, e), this suggests that rod photoreceptors may have similar asymmetries in responding to positive and negative intensity changes as cone photoreceptors. This could in principle explain why the contrast representations remain unaltered at different ambient light levels, even when retinal circuits can alter their responses considerably (Tikidji-Hamburyan et al., 2015). Nonetheless, retinal pathways downstream of photoreceptors can also be involved in modifying the response to different contrast levels (Freed, 2017; Zaghloul et al., 2003).

Our finding that ‘contrast’ means different quantities for ON and OFF RGCs has several implications in vision science. By default, contrast is often considered to be a symmetric quantity with respect to positive and negative luminance changes, meaning that ON and OFF RGCs encode equal but opposite luminance changes. Most visual studies build on this assumption to design contrast-balanced stimuli in order to stimulate the ON and OFF pathways equally. Even more importantly, stimuli should be designed in a way that all ON RGCS, irrespective of their spatial location on the retina, are activated equally, and the same would be expected for OFF RGCs. For example, according to our data, if one wanted to activate all OFF RGCs equally by a “flash” (dark flash) on top of a non-homogeneous scene (e.g. a natural image, where the starting intensity is different for RGCs distributed across the retina), then a constant number should be *subtracted* from each pixel in the scene. To activate all ON RGCs equally, all pixel-intensities of that scene should be changed by first *multiplying* by a constant factor (representing the fact that ON RGCs partially behave according to ratio contrast), and then *adding* a constant value. Our findings can therefore lead to a more accurate design of visual stimulation paradigms. Overall, our results describe what contrast means for ON and OFF RGCs which is crucial in understanding natural vision, given that the retinal sensitivity is governed by the magnitude and polarity of frequent intensity changes resulting from eye movements such as saccades (Idrees et al., 2020a, 2020b). Our results also demonstrate a novel asymmetry between ON and OFF pathways in the retina, which is consistent with the notion that ON and OFF pathways carry qualitatively different types of information (Chichilnisky and Kalmar, 2002; Pandarinath et al., 2010).

## Acknowledgements

This research was supported by Deutsche Forschungsgemeinschaft (DFG) grant MU3792/1-1 to TAM. TAM also received support from the Tistou and Charlotte Kerstan Foundation. We thank Jithin Nambiar and Nasser Karmali for assessing the quality of units sorted by the automated spike sorter and applying the necessary post-hoc steps to obtain high-quality units. We thank Timm Schubert and Felix Franke for comments on the manuscript.

## Author contributions

TAM conceptualized the study; SI and TAM designed the overall study; SI performed the retina electrophysiology experiments; SI analyzed the data with supervision from TAM. SI and TAM interpreted the data and wrote the manuscript.

## Data availability

All data presented in this paper are stored and archived on secure institute computers and are available upon reasonable request.

## Methods

### Animals

Mouse ex vivo retina electrophysiology experiments were performed in Tübingen, in accordance with German and European regulations, and animal experiments were approved by the Regierungspräsidium Tübingen.

We used 3 retinae from 2 male and 1 female *PV-Cre x Thy-S-Y* mice (*B6;129P2-Pvalb^tm1(cre)Arbr^/J × C57BL/6-tg (ThystopYFPJS))*, 5-9 months old, which are functionally wild type (Farrow et al., 2013; Münch et al., 2009; Tikidji-Hamburyan et al., 2015). We housed mice on a 12/12 h light/dark cycle, in ambient temperatures between 20-22 °C and humidity levels of 40%.

### Procedure and laboratory setup

Mice were dark adapted for 4-16 h before experiments. We then sacrificed them under dim red light, removed the eyes, and placed eyecups in Ringer solution (in mM: 110 NaCl, 2.5 KCl, 1 CaCl_2_, 1.6 MgCl_2_, 10 D-glucose, and 22 NaHCO_3_) bubbled with 5% CO_2_ and 95% O_2_. We removed the retina from the pigment epithelium and sclera while in Ringer solution.

We recorded retinal ganglion cell (RGC) activity using the MaxOne high-density multielectrode array (MEA) system (Müller et al., 2015) (Maxwell Biosystems, Basel, Switzerland). The MaxOne MEA featured 26,400 metal electrodes with center-to-center spacing of 17.5 μm in a grid-like arrangement over an area of 3.85 x 2.1 mm. Up to 1024 electrodes could be selected for simultaneous recordings. For each experiment, a piece of isolated retina covering almost the entire electrode array was cut and placed RGC-side down in the recording chamber. We achieved good electrode contact by applying pressure on the photoreceptor side of the retina by carefully lowering a transparent permeable membrane (Corning Transwell polyester membrane, 10 μm thick, 0.4 μm pore diameter) with the aid of a micromanipulator. The membrane was drilled with 200 μm holes, with center-center distance of 400 μm, in a regular hexagonal arrangement, to improve access of the Ringer solution to the retina. We superfused the tissue with Ringer solution at 30-34 °C during recordings, and we recorded extracellular activity at 20 kHz using FPGA signal processing hardware. Data were acquired using MaxLab software provided by Maxwell Biosystems, Basel, Switzerland.

We presented light stimuli to the retinal piece that was placed on the MEA using a DLP projector running at 60 Hz (Lightcrafter 4500 from EKB Technologies Ltd.) with internal red, green and blue light-emitting diodes. The projector had a resolution of 1280 x 800 pixels, extending 3.072 x 1.92 mm on the retinal surface. We focused images onto the photoreceptors using a 5x objective (illumination from above). The light path contained a shutter and two motorized filter wheels with a set of neutral density (ND) filters (Thorlabs NE10B-A to NE50B-A), having optical densities from 1 (ND1) to 5 (ND5). The filters allowed us to adjust the absolute light level of the stimulation.

We measured the spectral intensity profile (in μW cm^-2^ nm^-1^) of our light stimuli with a calibrated USB2000+ spectrophotometer (Ocean Optics) and converted the physical intensity into a biological equivalent of photoisomerizations per rod photoreceptor per second (R*rod^-1^s^-1^), as described before (Tikidji-Hamburyan et al., 2015). Light intensities of the projector output covered a range of 3 log units (i.e. 1000-fold difference between black and white pixels, over an 8-bit range). We used the Lightcrafter projector in pattern sequence mode. In this mode, the projector output is linear. Absolute light intensities at the mesopic level ranged between 9 R*rod^-1^s^-1^ for our darkest stimuli to 8169 R*rod^-1^s^-1^ for our brightest stimuli. At the scotopic level, the intensities were 100 times dimmer and at the higher mesopic level the intensities were 10 times brighter.

### Visual stimuli

Our visual stimulus consisted of uniform full-field steps. Each step consisted of an *Intensity_before_ → Intensity_after_* step where the display intensity was maintained at the value *Intensity_before_* for 4 seconds, followed by an instantaneous change to *Intensity_after_*. After 1 second at *Intensity_after_*, we switched to the next *Intensity_before_* value. A single trial consisted of 39 successive *Intensity_before_ → Intensity_after_* steps. The *Intensity_before_* values ranged from 2157 to 4304 R*rod^-1^s^-1^ at mesopic light level. *Intensity_after_* values ranged from 9 to 8169 R*rod^-1^s^-1^. In total there were 368 such steps that induced light increments and 256 steps that induced light decrements across the retina. These 624 *Intensity_before_ → Intensity_after_* steps were distributed pseudo-randomly across 16 trials. The 624 dots in Fig. 2 illustrate all these steps. The batch of 16 trials was repeated at least 4 or 5 times at a single ambient light regime.

### Data analysis

#### MEA recordings preprocessing

For high-density MEA recordings, we performed spike sorting by an offline automatic algorithm (Diggelmann et al., 2018). The sorted units were curated with a custom developed tool, the UnitBrowser (Idrees et al., 2016). We judged the quality of all units using inter-spike intervals and spike shape variation. Low quality units, such as ones with high inter-spike intervals, missing spikes, or contamination, were discarded. All spike rate analyses were based on spike times of individual units. In total, we extracted and analyzed 243 high quality units after the spike sorting (referred to as RGCs from now on). We converted spike times to estimates of spike rate by convolving these times with a Gaussian of *σ* = 10 ms standard deviation and amplitude 0.25 *σ*^−1^e^1/2^

#### Peak responses to *Intensity_before_ → Intensity_after_* steps

For each RGC, we calculated a response to each *Intensity_before_ → Intensity_after_* step by averaging the RGC’s spike rate to all repetitions of that step. RGCs that had stronger responses to light increments were classified as ON RGCs, and RGCs that had stronger responses to light decrements were classified as OFF RGCs. ON RGCs were then further analyzed using only steps with *Intensity_before_ < Intensity_after_* (light increments), and OFF RGCs were further analyzed using only steps with *Intensity_before_ > Intensity_after_* (light decrements).

For all RGCs, we then calculated a peak response to each *Intensity_before_ → Intensity_after_* step as the maximum response within 400 ms from the time of step (“response strength”). We discarded the response to a particular *Intensity_before_ → Intensity_after_* step if the peak response was within noise levels, i.e. within 4 standard deviations of the background response (1000 ms prior to the step). For each RGC, we then normalized the peak response to every *Intensity_before_ → Intensity_after_* step by the median peak response across all steps (we did not normalize by the maximal peak so that potential saturation of responses for stronger stimuli would not influence the overall shape of the stimulus-response relationship).

We then generated a generic ON RGC by taking the median across all ON RGCs for each before/after combination. Finally, we normalized the resulting responses by the strongest response (Fig. 2a, black dots). We fitted a second-order polynomial to the resulting responses in order to obtain responses to a continuous range of *Intensity_before_ → Intensity_after_* combinations. Correspondingly, we generated a generic OFF RGC. The responses of these generic ON and OFF RGCs (Fig. 2a) were used for all analysis stated in the results section.

To obtain the time-varying response traces (Fig. 2c), we repeated the above procedure on the spike rates and not the peak responses.

All data analyses were performed in MATLAB (The MathWorks Inc).

